# Cell Population Effects in a Mouse Tauopathy Model Identified by Single Cell Sequencing

**DOI:** 10.1101/771501

**Authors:** Véronique Lisi, Gabriel Luna, Angeliki Apostolaki, Michel Giroux, Kenneth S Kosik

## Abstract

Neurodegenerative disorders are complex multifactorial diseases that have poorly understood selective vulnerabilities among discrete cell populations. We performed single cell RNA sequencing of whole hippocampi from the rTg4510 mouse tauopathy model, which expresses a P301L *MAPT* mutation at two time points—before and after the onset of pathology. One population of neurons showed a robust size reduction in both the young and the old transgenic animals. Differential expression of genes expressed in this group of neurons suggested an enrichment in granule cell neurons. We identified genes that characterize this population of neurons using Pareto optimization of the specificity and precision of gene pairs for the population of interest. The resulting optimal marker genes were overwhelmingly associated with neuronal projections and their expression was enriched in the dentate gyrus suggesting that the rTg4510 mouse is a good model for Pick’s disease. This observation suggested that the tau mutation affects the population of neurons associated with neuronal projections even before overt tau inclusions can be observed. Out of the optimal pairs of genes identified as markers of the population of neurons of interest, we selected Purkinje cell protein 4 (*Pcp4+*) and Syntaxin binding protein 6 (*Stxbp6+*) for experimental validation. Single-molecule RNA fluorescence *in situ* hybridization confirmed preferential expression of these markers and localized them to the dentate gyrus.

## Introduction

Tauopathies are a class of neurodegenerative disorders that include, amongst others, Alzheimer’s disease, Pick’s disease, frontotemporal dementia, and progressive supranuclear palsy. These are all characterized by the misfolding and accumulation of tau protein. Key molecular and cellular mechanisms of the disease, such as selective cell vulnerability, remain poorly understood. Single-cell RNA sequencing (scRNA-seq) offers a means to determine cellular heterogeneity by profiling thousands of individual cells (Lake et al. 2016; Habib et al. 2017; Zhong et al. 2018; Lake et al. 2018; Macosko et al. 2015) in order to detect changes in transcriptionally similar cell populations. Comprehensive characterizations of the cell types present in various mammalian organs such as the dorsal root ganglia (Nguyen et al. 2017; Usoskin et al. 2014), the brain (Zeisel et al. 2015; Tasic et al. 2016), the kidney (Park et al. 2018), and intestinal organoids (Grün et al. 2015) have been published. In the context of Alzheimer’s disease, single cell sequencing of immune cells has revealed the existence of a “disease-associated microglia” state in Alzheimer’s disease (Keren-shaul et al. 2017). In a larger study of Alzheimer’s brain tissue, an over-representation of female cells was present in AD subpopulations and myelination-related processes were perturbed in multiple cell types (Mathys et al. 2019).

We sought to assess cell population changes in the mouse tauopathy model, rTg4510, by profiling thousands of individual cells at two different time points. rTg4510 animals express a mutant form of human Tau (P301L) and exhibit behavioral deficits, neuronal loss, microgliosis and neurofibrillary tangles (Ramsden et al. 2005). Unbiased clustering of single neuron transcriptional profiles identified a sub-population of neurons that was depleted in the rTg4510 animals when compared to their littermate controls at both a young age, i.e. before detection of Tau pathology, and at an older age when the tauopathy was manifest. This cluster of neurons could not be assigned a neuronal identity based on marker genes or gene ontology. To gain insight into the identity of this cluster, we identified genes that were uniquely expressed together in the neurons of the cluster of interest, but not co-expressed in the neurons of the other clusters. Optimal gene pairs, according to a Pareto optimization based on sensitivity and precision criteria were then identified. We validated one of the gene pairs by single molecule fluorescence *in situ* hybridization. The set of optimal marker genes identified in this manner was associated with neuronal projections.

## Results

To study the effect of Tau^P301L^ overexpression on cellular identity we performed single-cell RNA sequencing using the Dropseq method (Macosko et al. 2015) on cells isolated from the hippocampi of three rTg4510 and four control animals at a young age (4-6 weeks old), and four rTg4510 and 5 control animals at an older age (32-40 weeks old). At the younger time point, behavioral deficits were observable before the appearance of Tau pathology, as detected by MC1 staining (Hernandez et al. 2019), suggesting that Tau overexpression impacted cells before the deposition of fibrillar aggregates could be observed. We sequenced 20,900 cells from which 19,539 cells remained after quality control (see Methods), distributed as follows: 5616 cells from the young control animals, 3624 from the young rTg4510 animals, 4776 from the old control animals and 5523 from the old rTg4510 animals.

### A population of neurons is depleted in the transgenic animals

To identify neuronal populations that differ between the rTg4510 and control animals, we first classified cells with a set of published markers for cell types of the nervous system (see Methods). Of the 19,539 cells remaining after quality control, 7600 were identified as neurons, 3968 as oligodendrocytes, 3882 as astrocytes, 3437 as microglia and 652 cells were of a non-specific glial identity (Fig. 1). Of the 7600 neurons, 2596 came from the young control animal, 1406 from the young transgenic, 1826 from the old control and 1772 from the old transgenic. We focused on the neuronal population and removed from the analysis those genes that were not expressed in at least ten neurons. To identify neuronal types that differed between the rTg4510 animals and the littermate controls, we performed dimensionality reduction and clustering using the shared nearest neighbor technique implemented in the Seurat package version 2 (Butler et al. 2018). In doing so, 26 neuronal clusters were identified (Fig 2A). Of these, 7 clusters were major clusters, together comprising over 50% of the all neurons (Fig 2A, clusters 0 to 6 and Supplementary Table I). We evaluated the robustness of these seven major clusters on simulated data constructed by adding normally distributed noise to the original count matrix. We then renormalized and rescaled the data and re-clustered the cells. We computed the Jaccard index for each of the new clusters (see Methods). Doing so, four of the abundant clusters were robustly identified in the simulations as evidenced by a mean Jaccard index greater than 0.5 (Fig 2B, clusters 0, 1, 2, and 4).

**Figure 1.**
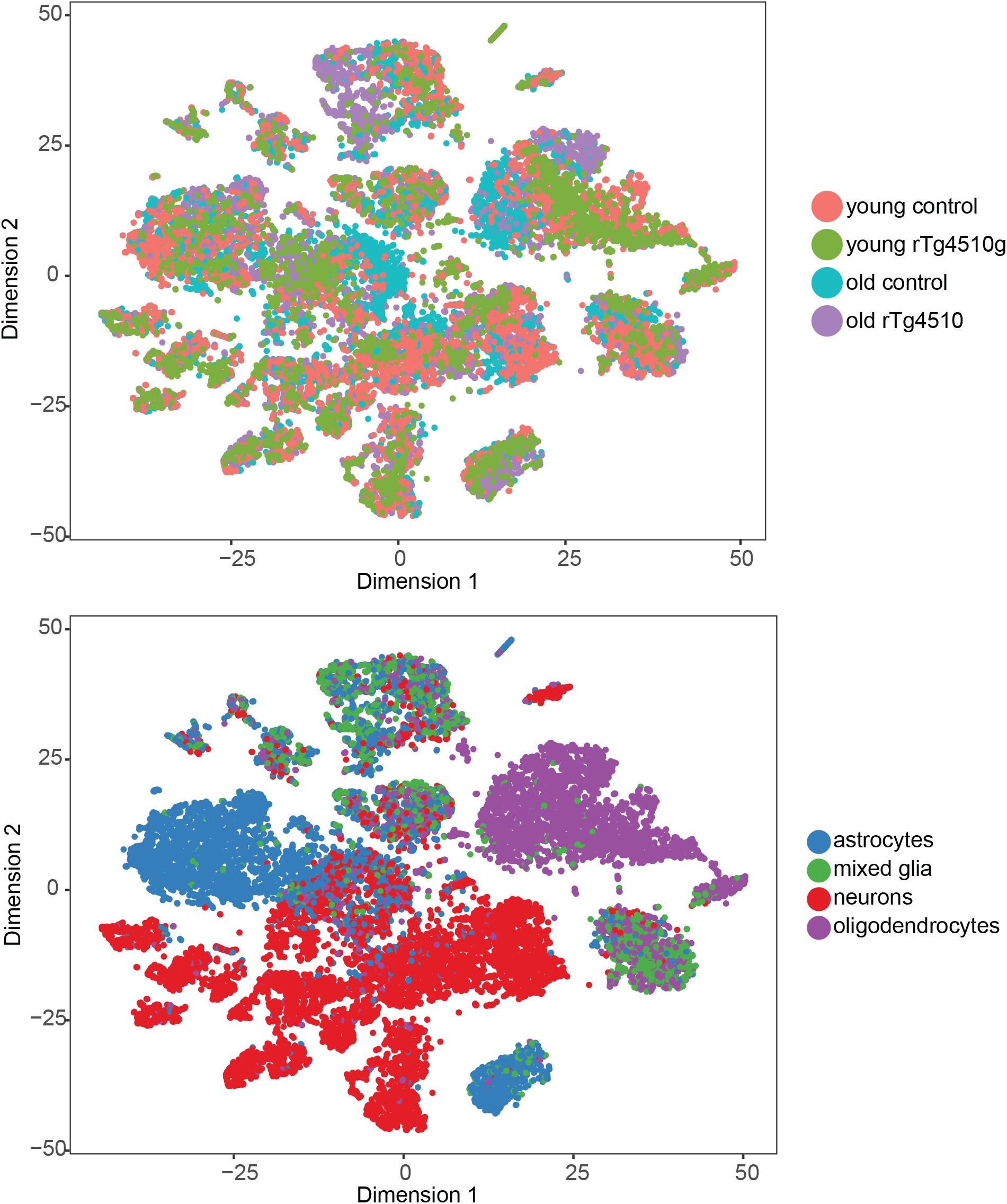
tSNE embedding of the data. Expression data was reduced in dimension by PCA and then embedded in two dimension using tSNE. Cells are color coded according to their genotype/age (top panel) or their assigned identity (bottom panel). Cells from each genotype/age is found in every cell type assignements and cells of the same type cluster together, as expected. Further analysis focused on the cells assigned neuronal identity.

**Figure 2.**
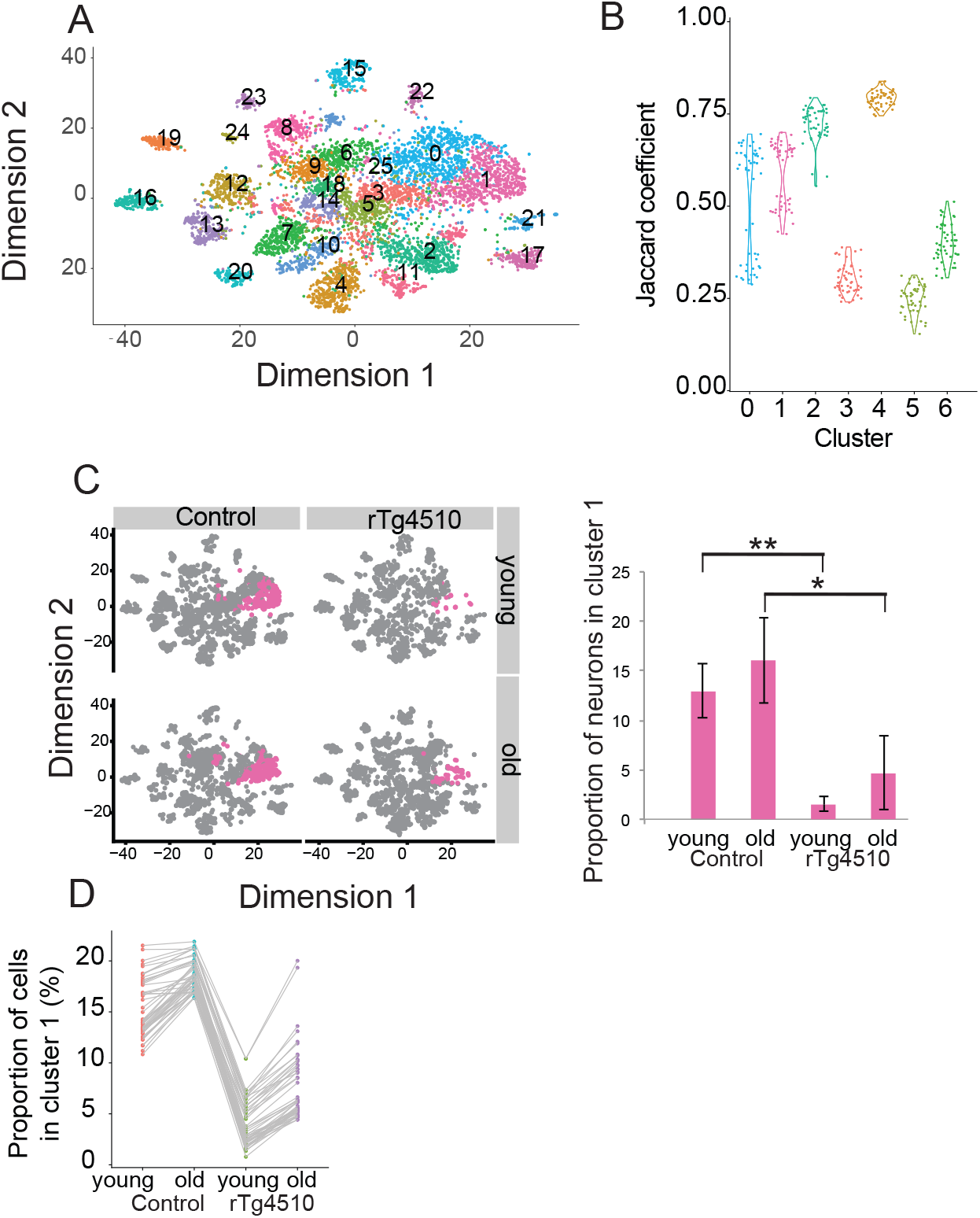
Identification of neuronal clusters whose abundance differs between the rTg4510 and the control animals. A. tSNE representation of the neurons after dimensionality reduction and cluster identification. Clusters labeled 0 to 6 comprise over 50% of all the neurons. B. Jaccard coefficient of the simulations for the 7 abundant clusters. Clusters 0,1,2, and 4 are robustly identified in the simulations with a mean jaccard coefficient greater than 0.5. C. tSNE representation of the neurons as in A but split according to genotype and age and highlighting only cluster 1. The proportion of neurons in cluster 1 is shown on the right. The rTg4510 animals have a smaller proportion of neurons in cluster 1 than the control animals. [The mean proportion per animal is presented, the error bar represents the SEM; *t.test pvalue < 0.05; **t.test pvalue < 0.01]. D. Proportion of neurons in cluster 1 for each sample type in the simulated dataset. Each gray line represents 1 simulation. For every simulation, the proportion of neurons in cluster 1 is smaller in the rTg4510 samples.

We focused our attention on cluster 1 that underwent a highly robust change between the control animals and both the young and old rTg4510 animals. In both the young and old control animals the proportion of neurons in cluster 1 was lower in the rTg4510 animals than in the control animals in each of the animals we analyzed (Fig. 2C – right panel). We confirmed that the cell population differences were robust to noise by generating simulated datasets with the addition of normally distributed noise to the single cell sequencing counts matrix. In every simulation generated, the proportion of neurons in cluster 1 was lower in the rTg4510 samples than in the control samples (Fig. 2D). We further confirmed that the cell proportion differences were not an artifact of the clustering method by applying a second clustering method from Monocle2, density peaks clustering (Qiu et al. 2017), which resulted in a similar decrease in a cluster of neurons in the rTg4510 animals (Fig. S1 and supplementary information).

Using the same method to analyze the microglia, we found a sub-cluster of cells that was almost exclusively present in the old rTg4510 animals. This cluster expressed genes related to microglial activation such as Aif1, Cd68, Il1b, Ptprc (aka Cd45) and Tnf. Thus, our method confirmed the higher abundance of activated microglia in the old rTg4510 animals as previously reported (Lee et al. 2010).

The reduced cluster 1 abundance in the rTg4510 animals was observed even before tau inclusions were detectable. Therefore, the gene expression profiles of that cluster could potentially be informative of the earliest molecular processes underlying tau pathology. We thus sought to identify the genes that define cluster 1. We attempted to assign a specific neuronal cell identity to this cluster by inspecting the expression of known marker genes for known neuronal cell types. This approach failed to identify a specific recognized neuronal cell type indicative of the cluster. Therefore, we sought to characterize this cluster by identifying sets of genes that, when taken together, precisely and specifically describe the neurons of cluster 1. Differential gene expression analysis between the neurons of cluster 1 and the neurons of the other clusters can provide information as to the identity of the neurons in cluster 1. This approach however is inaccurate when the number of cells being compared is imbalanced; nor does it consider the potential synergy of genes as markers of a group of cells i-e that two genes with modest specificity and precision for a cellular population on their own are highly specific and precise for a population of cells when combined. Using these two parameters — specificity and precision — we devised a method to identify optimal sets of genes that described the neurons of cluster 1. Briefly, for each gene, we compute two measures that together, define how useful the marker gene is: the precision and sensitivity. A perfect marker would identify all of the neurons of cluster 1 (sensitivity of 1) and only the neurons of cluster 1 (precision of 1) (see Methods). We plotted each gene as precision *vs* sensitivity and use a Pareto optimization to find sets of optimal markers, i.e. those genes for which no other gene has both a greater precision and sensitivity. Doing so, we identified nine optimal single markers (Fig. S2). To identify optimal pairs of markers, we repeated the procedure using a greedy approach, testing only the pairs that included one of the optimal single markers. This resulted in 19 unique pairs of optimal markers (Fig. 3A). As expected, pairs of genes used as markers had increased precision and sensitivity over single genes. The pairs of optimal markers making up the pareto front contained 18 unique genes. These genes were overwhelmingly associated with neuronal projections; 10 of these 18 genes are known to be implicated in neuron projection. These include Gap43, Calm1, Ppp3ca, Gria2, Ncdn, Pcp4, Nrgn, Olfm1, Plk5 and Penk. Of the other 8 genes, 4 were associated with other neuronal related functions or localization: Scg2 (neuron components), Stxbp6 (SNARE complex), C1ql3 (synapse organization) Trpc6 (neuron differentiation). Finally, 4 genes had either no known function (Snhg11) or had main functions not specifically related to neuronal function: Ppp1r1a and Rasl10a (signal transduction), Rbp4 (glucose and retinoid process)

**Figure 3.**
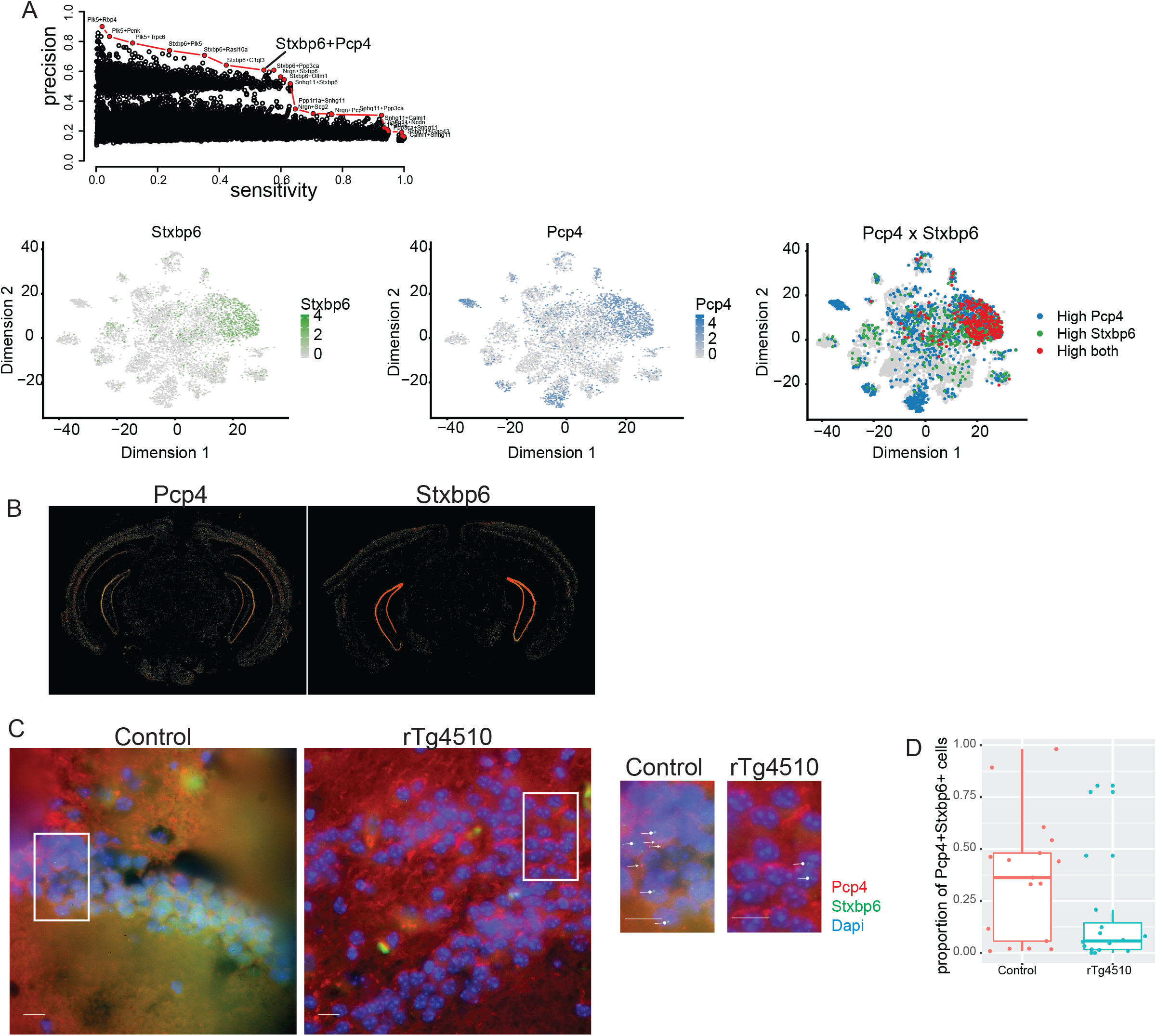
Characterization of cluster 1. A. Sensitivity and precision of the marker pairs. The optimal pairs of markers are identified by a Pareto optimization, highlighted in red, and labeled. There are 22 such pairs including the Pcp4/Stxbp6 pair (top panel). Both Stxbp6 and Pcp4 are enriched in the cells of cluster 1 and expressed by cells from other clusters. Double positive cells are found almost exclusively in the cells of cluster 1 (bottom right panel) B. In situ hybridization data from the Allen Brain Institute mouse atlas for adult (P19) animals showing that both Pcp4 and Stxbp6 are expressed in the dentate gyrus. C. Representative images of the single molecule fluorescence in situ hybridization. Inset are scaled at the right of the image. Round arrowheads highlight Stxb6 signal whereas triangular arrowheads highlight Pcp4 signal. Scale bar: 10um. D. Quantification of the smFISH images. The proportion of cells positive for both Stxbp6 and Pcp4 is higher in the control samples than in the rTg4510.

To validate the computational method and better characterize the expression of these optimal markers in neurons of the brain, we selected a marker pair for experimental validation. Given that each of the pairs are optimal, we devised another metric for selecting the marker pair for validation. We computed, for each pair, the proportion of double-positive cells in our single cell transcriptome data for each sample type. We also computed the ratio of this proportion between the old rTg4510 and old control samples and between the young rTg4510 and young control samples in our single cell data (Table I). We selected the marker pair with the highest fold change between the number of double-positive cells in the old rTg4510 and the old control samples, but limited our selection to the marker pairs for which at least 5% of the cells were double positive in each of the samples (old rTg4510 and old control) (Table I). Based on these criteria, we selected the marker pair Stxbp6 (Syntaxin binding protein 6, also known as amysin) and Pcp4 (Purkinje cell protein 4, also known as Pep19) for further characterization. In our single cell data, 14% of the old control cells and 5% of the old rTg4510 are double positive for Stxbp6 and Pcp4. These proportions of cells were also observed in the young animals; 16% of the young control cells and 5% of the young rTg4510 cells are double positive for Stxbp6 and Pcp4. In general, the proportion of double positive cells for each marker was similar when comparing the same genotype across time (young vs old rTg4510 or control) but differed when comparing the genotypes at the same time point (old control vs rTg4510) (Table I). Importantly, while the number of double-positive cells differed between the old rTg4510 mice and their littermate controls, the expression level of either gene was not significantly different between the two genotypes at either time point (Fig. 3A). Therefore, this metric identified exclusively cell population differences without an effect on the transcript level in the experimental and control cell populations.

**Table I.**
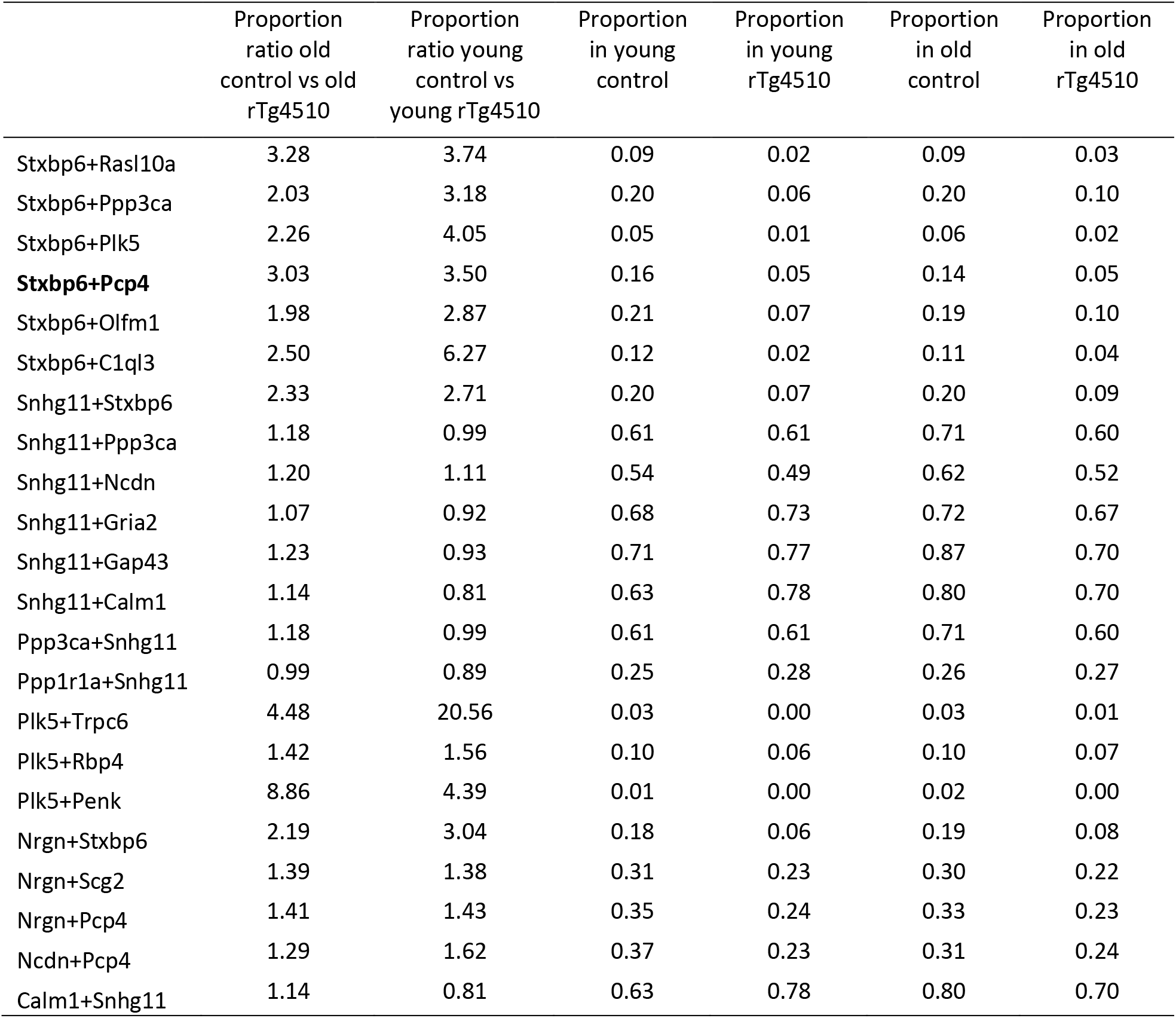
Predicted proportion and fold change of the optimal marker pairs identified. Stxbp6+Pcp4 (in bold) were selected for experimental validation because of the combined relatively high expected proportion in both strains at both timepoints and the expected high proportion fold change at both time points.

We next looked for brain regions which expressed both Stxbp6 and Pcp4 by interrogating the Allen Brain Institute mouse atlas (Lein et al. 2007). Both genes are enriched in the cells of the dentate gyrus (Fig. 3B). This enrichment in the dentate gyrus was also observed for at least one of the genes in every gene pair identified as an optimal marker pair by Pareto optimization, strongly suggesting that the cluster of neurons of interest was physically located in this region. We compared these (314) genes to the Mammalian Adult Neurogenesis Gene Ontology (Mango) (Overall, Paszkowski-Rogacz, and Kempermann 2012) and computed the enrichment of each of the terms in the ontology. This analysis identified the ontology term “Granule cell neuron” consistent with enrichment of this population in the dentate gyrus.

To validate the existence of the population of neurons found by single cell sequencing, we performed single molecule RNA fluorescence *in situ* hybridization (smFISH) of both control and rTg4510 mouse cryosections. We concentrated our analysis on the hippocampus, specifically the dentate granule cells which were enriched with the two marker genes. We quantified the proportion of double positive (Stxbp6^+^/Pcp4^+^) cells in each of the samples (Fig. 3C). As predicted by the computational approach based on the single cell transcriptomes, a greater proportion of double positive cells were found in the control sections than in the rTg4510 sections (Fig. 3D).

Given that the rTg4510 mouse model requires the breeding of two strains each carrying a single transgene, we wanted to ensure that our results were only observed in the presence of both transgenes. The P301L MAPT mutation is under a tetO promoter that is activated in the brain when the animals are bred to CamK2a-tTa animals. We performed single cell RNA sequencing of old animals carrying either of the two alleles. The single transgene animals are each on a different background and are thus on a different genetic background than that of the rTg4510 and their littermate controls. A combined analysis of all of these animals is therefore not informative because the genetic background dominates the analysis. Analyzing the single transgene independently, the cluster containing the Stxbp6+/Pcp4+ neurons was composed of 10% of the P301L neurons and 24% of the rTta neurons showing that the observed low abundance of Stxbp6+/Pcp4+ neurons of cluster 1 in the rTg4510 animals was not driven by either single transgene alone.

## Discussion

Using single-cell sequencing, we found a population of neurons that is less abundant in the rTg4510 animals than in their littermate controls and developed a computational method to identify markers of this population. This method identifies combinations of genes that together are optimally precise and sensitive towards the population of cells under study. This method can be applied to the characterization of any cluster of cells identified by single cell sequencing. Applying our method to the characterization of cluster 1, we identified Pcp4 and Stxbp6 as one pair of markers that define this cluster. Pcp4 expression has been shown to be decreased in the brain of patients with Alzheimer’s disease and Huntington’s disease (Utal et al. 1998) and in the brain of alcoholic patients (Iwamoto et al. 2004). Out of the optimal markers for the cells of cluster 1 identified by the pareto front, Gap43 (de la Monte, Ng, and Hsu 1995), Calm1 (Esteras et al. 2012), Nrgn (Thorsell et al. 2010) and Trpc6 (Lessard et al. 2005) have also been reported as aberrantly expressed or regulated in Alzheimer’s disease.

Adult transgenic mice overexpressing Pcp4 show earlier maturation of neurons compared to wild type littermate controls (Mouton-Liger et al. 2013). This point is of interest because we also observed a parallel increased abundance of another neuronal population designated cluster 0 in the rTg4510 animals. Comparing these two populations of neurons suggested that the neurons of cluster 1, those that are less abundant in the rTg4510 animals, are more mature than those of cluster 0. Together, these results associate the behavioral deficits observed in the early stages of rTg4510 progression with a neuronal maturation deficit or the emergence of a precursor population. We confirmed the finding of Pcp4 and Stxbp6 as marker genes for the reduced population by single molecule RNA FISH. This localization suggested a resemblance of rTg4510 mouse model to Pick’s disease, a tauopathy with selective vulnerability of dentate granule cells.

The two marker genes for the cluster that decreased with disease progression are exclusively co-detected in the dentate granule cells. The dentate gyrus is a major area in which neurogenesis takes place in the adult mammalian brain (Eriksson et al. 1998) even at older ages (Kempermann, Kuhn, and Gage 1997) and is thought to play a role in hippocampal-dependent learning and memory (Gould et al. 1999; Shors et al. 2001; Snyder et al. 2005) The effects on neurogenesis (reviewed in (Rodríguez and Verkhratsky 2011)) and cell cycle re-entry (reviewed in (Yang and Herrup 2007)) in Alzheimer animal models are complex. Most of these findings pertain to models of beta-amyloid deposition. Much less is known about neurogenesis in the tauopathies. Studies of this topic must address the balance between proliferation and differentiation. This balance appears to shift toward proliferative cells in the Alzheimer brain (Stopa et al. 1990; Hock et al. 2000). Compared to controls, Alzheimer patients’ brains increase the expression of doublecortin, polysialylated nerve cell adhesion molecule, neurogenic differentiation factor and TUC-4 (Jin et al. 2004). Doublecortin and TUC-4 are associated with neurons in the neuro-proliferative zone of the dentate gyrus suggesting that neurogenesis is increased in the Alzheimer hippocampus, where it may give rise to cells that replace neurons lost in the disease (Jin et al. 2004). The localization of tau pathology to these regions suggests that the tauopathy may be responsible for effects of neurogenesis, which would result in the more immature profile we observed here. Furthermore, the entire cluster of genes in the Pareto front directed toward axonal projections is consistent with another P301L tau over-expressing transgenic mouse in which tau occupies the terminal zone of performant pathway (Pooler et al. 2013). We do note the caveat in a recent report the observed phenotype in rTg4510 animals could be the result of the disruption of six endogenous genes caused by the insertion of the Tau and tTa allele (Gamache et al. 2019). However, none of these genes were identified as markers of the population of neurons we identified.

## Methods

### Animals

#### Tissue harvesting

Female rTg4510 that express Tau^P301L^ and rTTA were euthanized with CO2 at young (4-6 weeks old) and at old age (32-39 weeks old). The hippocampus was dissected out, digested with papain and mechanically triturated. Cell suspension was layered on 4% BSA and spun at 300g for 10 minutes to remove debris. The pellet was resuspended in PBS and processed for dropseq as described (Macosko et al. 2015). Mice from both strains were processed together from dissection to droplet generation and cDNA generation. Once two animals from each time points were processed to cDNA generation, pre-amplification, tagmentation, final library preparation and sequencing were done with all samples in parallel. This whole process was repeated for the other animals. All animal experiments were performed in accordance to the University of California Santa Barbara Institutional Animal Care and Use Committee approved protocols.

#### Immunocytochemistry

Animals were transcardially perfused using 4% paraformaldehyde in 0.1 M sodium cacodylate (Electron Microscopy Sciences, Hatfield, PA) for 15 min at room temperature. Brains were then dissected and immersion fixed for 4 h at 4°C. Samples were rinsed 5 x 5 min in phosphate buffered saline (PBS; pH 7.4) and then cryoprotected in a graded sucrose solution, samples were subsequently sectioned at 10 µm using a cryostat (Leica, Lumberton, NJ). Sections were immersed in normal donkey serum 1:20 in PBS containing 0.5% BSA, 0.1% Triton-X 100, and 0.1% sodium azide (PBTA) for 30 mins at room temperature followed by immersion in anti-MC1 diluted in PBTA for 1 h (Peter Davies, Feinstein Institute for Medical Research, Manhasset, NY; mouse monoclonal, 1:200). Next, sections were rinsed in PBTA 5 x 5 min, 1 x 1 h in PBTA and then placed in secondary antibodies donkey anti-mouse 488 as well as the nuclear stain Hoechst (Molecular Probes, Eugene Oregon; 1: 5000) for 1 h at room temperature. Finally, secondary antibodies were rinsed in PBTA and mounted using Vectashield (Vector Laboratories Inc., Burlingame, CA) on a glass slide and sealed under an 18 x 18 #0 micro coverslip (Thomas Scientific; Swedesboro, NJ) using nail polish.

For single molecule fluorescence in situ hybridization, probes were designed using Stellaris probe designer. Pcp4 probes were labeled with Quasar-670 and Stxbp6 probes with Quasar-570. After cryosectioning, samples were briefly thawed and permeabilized in cold 70% EtOH at 4°C for at least 2 hours and up to overnight. Samples were rehydrated in 2X SSC for 3-5 minutes and treated with proteinase K solution in 2X SSX (100ng/ml) for 5 minutes at room temperature. Proteinase K treatment was stopped by rinsing twice with glycine (2mg/ml) in 2X SSC and the glycine washed three times with 2X SSC. 2ul of stellaris probes were diluted in 100ul of Stellaris hybridization buffer (Biosearch technologies SMF-HB1-10). Samples were hybridized overnight at 37°C, washed once with Stellaris wash buffer A (Biosearch technologies SMF-WA1-60) containing Dapi for 30 minutes at 37°C and then once with Stellaris wash buffer A (Biosearch technologies SMF-WA1-60) containing 4’,6-diamidino-2-phenylindole (DAPI) for 30 mins. at 37°C and then once with Stellaris wash buffer B (Biosearch technologies SMF-WB1-20) for three minutes at room temperature. Samples were treated with TrueBlack (Biotium) according to manufacturer instructions, mounted using Vectashield (Vector laboratories, Burlingame, CA) and imaged the same day.

#### Image acquisition

Specimens were screened and imaged using an Olympus Fluoview 1000 laser scanning confocal microscope (Center Valley, PA) equipped with an argon 488 argon photodiode laser, and HeNe 543/633 photodiode lasers as well as precision motorized stage (Applied Scientific Instrumentation, Inc. Eugene, OR). Sections that spanned the hippocampal formation from anterior to posterior used for quantitative analysis. Mosaics were captured using an UPlanSApo 20x oil immersion lens, N.A. 0.85 at 1 µm intervals along the z-axis and a pixel array of 800 x 800 in the x-y axes.

For the single molecule fluorescence in situ hybridization, specimens were imaged on an Olympus IX71 with a 60X water immersion objective. Twelve to fifteen Z-stacks of 0.3um were taken. Images were maximum projected and the smFISH signal deconvolved using a LoG filter. Cells were segmented on the DAPI signal using a watershed transformation and removing small cells using the EBImage package (Huber et al. 2010). smFISH signal per cell was counted as described (Lyubimova et al. 2013). Three slides per genotype were imaged each with five to seven field of views. The average field of view contained 91 cells for a total of 1599 cells analyzed for the WT samples and 1413 for the rTg4510 samples. The proportion of double positive cells over all the segmented cells per field of view is reported.

### Data processing and computational methods

#### Single cell RNA sequencing data processing

BCL files were transformed to fastq files using Illumina’s bcl2fastq setting the short adapter reads mask to 18. Fastq files were then transformed to BAM files using picard tools. BAM files were processed as described (Macosko et al. 2015) and in the dropseq cookbook v1.2 to assign reads to cells based on UMI, remove low quality reads, trim the adapters, align the reads and extract the digital expression matrix. This yielded 20900 cells. From this digital expression matrix, we removed 38 cells whose total number of counts was less than 100. We also removed 1323 cells for which the total number of transcripts detected was outside 2 standard deviations of the mean.

#### Classification of cell types

To classify cells as either neurons, astrocytes, oligodendrocytes or non specific glia, we used a set of published cell type markers (Ko et al. 2013). For each cell, we computed a normalized score to each cell type as described (Macosko et al. 2015). We defined four patterns of gene expression: only the genes of the neuron type are expressed, only those of the astrocyte type, only those of the oligodendrocytes or both those of the astrocytes and oligodendrocytes, representing uncharacterized glia. We assigned cell type based on the maximum correlation to these four patterns.

The counts for the cells identified as neurons were normalized and scaled using the Seurat package, regressing out the sequencing run and using a negative binomial distribution to scale the data. Using this transformed dataset, PCA was performed and tSNE projection and clustering were performed using the first 30 dimensions of the PCA.

#### Simulations

We generated 50 new datasets by adding normally distributed noise (mean = 0, sd = 2) to each count and rounding to zero the resulting negative counts. Using the new count matrix, we repeated the normalization, scaling, dimensionality reduction and clustering of the cells. We computed, for each initial cluster, its Jaccard coefficient in the new dataset.

#### Marker identification

To identify sets of genes that define the cluster of interest, we computed, for each gene, the sensitivity of this gene for the neurons of cluster 1 and the precision of the identification using this gene. To this end, we first computed, for each gene in the dataset, if the gene is a positive of negative marker of the neurons of cluster 1 by computing the fold change of the expression in the neurons of cluster 1 over the expression of all the other neurons. If the gene was a positive marker of the cluster, we counted the number of neurons expressing this gene, setting the expression threshold at the 5^th^ percentile of expression of this gene in the neurons of cluster 1. The ratio of positive neurons found in cluster 1 over all the positive neurons was the precision of the “isolated” population of neurons. The ratio of how many of the neurons of cluster 1 were found by this gene over all the neurons of cluster 1 was the sensitivity of the gene. We treated negative markers similarly, selecting negative cells as the ones whose expression is below the 95^th^ percentile. Of all the genes scored, we select the optimal ones using a Pareto optimization. To identify pairs of optimal markers, we computed, for each pair containing at least one optimal marker (as identified above), its purity and proportion by sequentially “isolating” the neurons, first on the optimal marker, then on the secondary marker.

## Acknowledgments

The authors acknowledge the use of the Biological Nanostructures Laboratory within the California NanoSystems Institute, supported by the University of California, Santa Barbara and the University of California, Office of the President. V.L. was funded by a fellowship from the Canadian Institutes of Health Research number 201511MFE-358735-159561. This work was supported by the Dr. Miriam and Sheldon G. Adelson Medical Research Foundation, the Rainwater Charitable Foundation and NIH grant U54NS100717 (K.S.K.)

## Competing interests

No competing interests declared

## Supplementary information

### Supplementary method

We applied the density peaks clustering method (from the Monocle package) and identified a cluster of neurons less abundant in the rTg4510 animals which was characterized by the same genes as the method presented in the main text.

**Figure S1 – Related to Figure 2.**
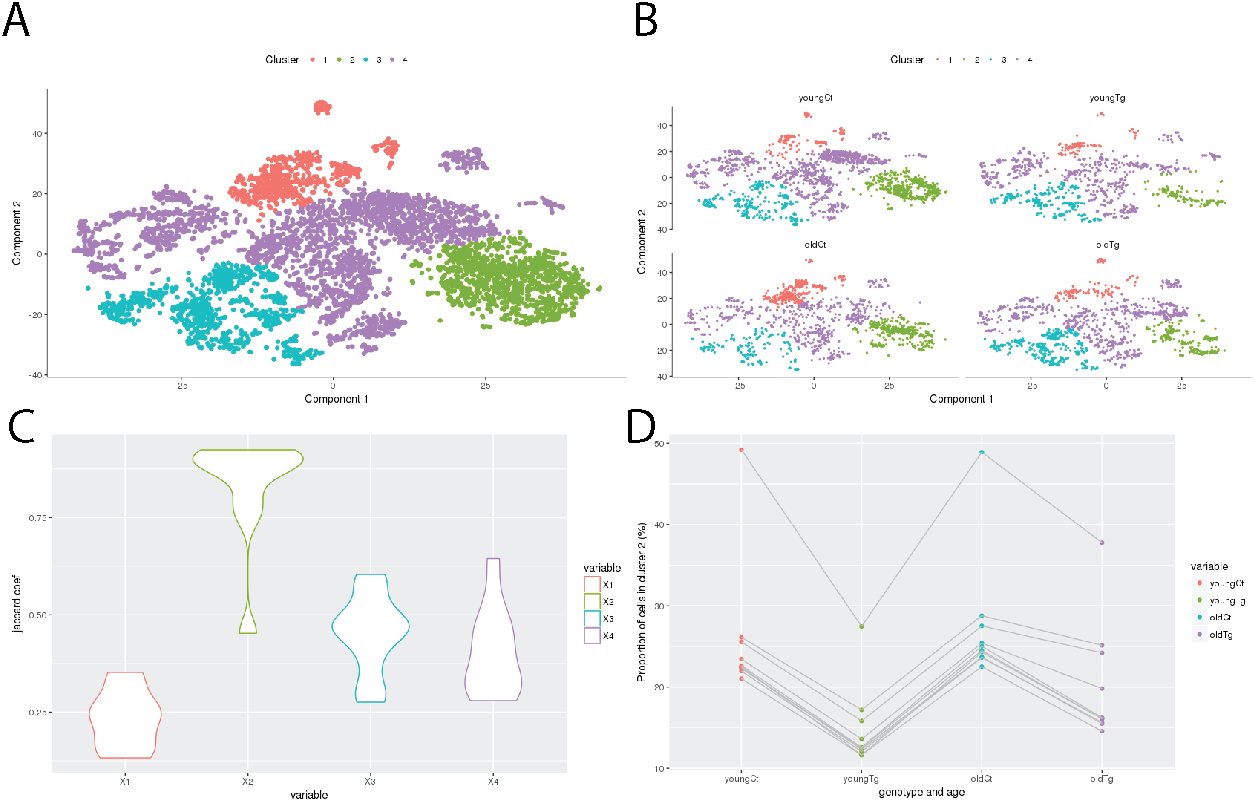
A) Neurons were clustered using density peaks which identifi clusters. B) Cluster 2 is less abundant in the rTg4510 animals. C) Cluster 2 is robust to noise addition (as described in Figure 2). D) In every simulation performed, cluster 2 is less abundant i the Tg animals than in the control animals.

**Figure S2.**
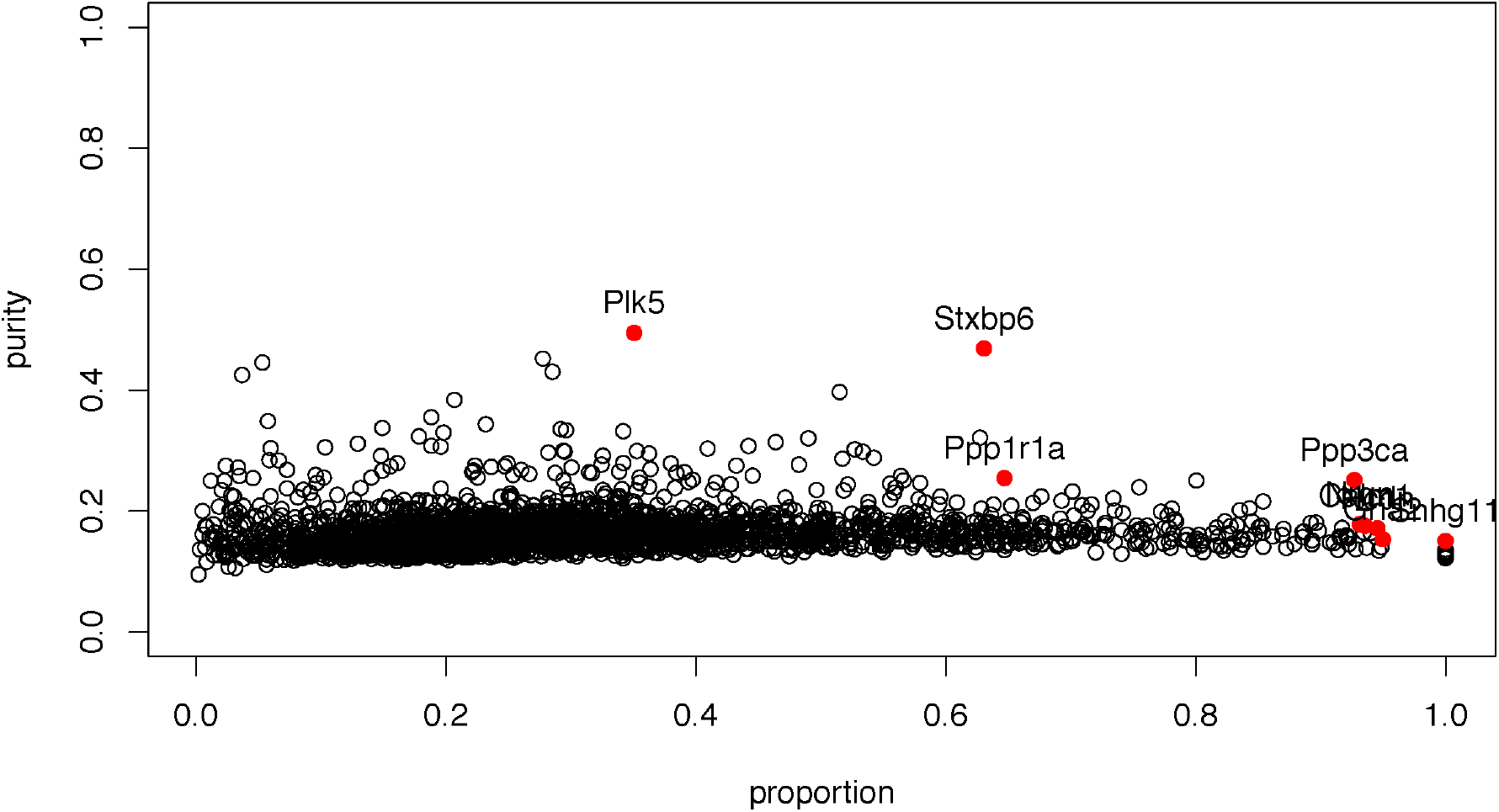
Purity vs propotion graph for the single genes with optimal genes highlighted.

**Supplementary Table I.** Number of neurons in each cluster.

| Cluster ID | Young |  | Old |  |
| --- | --- | --- | --- | --- |
|  | Control | rTg4510 | Control | rTg4510 |
| 0 | 259 | 142 | 103 | 317 |
| 1 | 305 | 17 | 330 | 78 |
| 2 | 167 | 90 | 71 | 190 |
| 3 | 336 | 28 | 59 | 59 |
| 4 | 114 | 105 | 76 | 124 |
| 5 | 181 | 98 | 52 | 78 |
| 6 | 31 | 46 | 241 | 54 |
| 7 | 134 | 61 | 93 | 74 |
| 8 | 113 | 87 | 57 | 85 |
| 9 | 32 | 99 | 91 | 82 |
| 10 | 105 | 48 | 61 | 72 |
| 11 | 68 | 87 | 18 | 103 |
| 12 | 92 | 74 | 48 | 59 |
| 13 | 83 | 68 | 33 | 59 |
| 14 | 90 | 22 | 62 | 46 |
| 15 | 53 | 39 | 68 | 59 |
| 16 | 75 | 52 | 32 | 42 |
| 17 | 69 | 27 | 64 | 32 |
| 18 | 21 | 5 | 129 | 15 |
| 19 | 66 | 28 | 20 | 47 |
| 20 | 47 | 52 | 21 | 16 |
| 21 | 33 | 25 | 29 | 22 |
| 22 | 40 | 59 | 5 | 4 |
| 23 | 33 | 33 | 12 | 15 |
| 24 | 6 | 7 | 15 | 27 |
| 25 | 4 | 3 | 1 | 5 |

